# Distinct Multimodal Imaging Correlates of Depression in Middle-Aged Adults With and Without a Family History of Alzheimer’s Disease

**DOI:** 10.64898/2026.04.13.717731

**Authors:** Barbara Duarte-Abritta, Carolina Abulafia, Leticia Fiorentini, Gustavo Tafet, Luis Ignacio Brusco, Aki Tsuchiyagaito, Sanjay J. Mathew, Mirta F. Villarreal, Salvador M. Guinjoan

**Affiliations:** Laboratory of Chronophysiology, Institute for Biomedical Research (BIOMED), Pontifical Catholic University of Argentina (UCA) and CONICET, Buenos Aires, Argentina; Consejo Nacional de Investigaciones Científicas y Tecnológicas (CONICET), Argentina; Fundación para la Lucha contra las Enfermedades Neurológicas de la Infancia (FLENI), Buenos Aires, Argentina; Department of Psychiatry and Mental Health, University of Buenos Aires School of Medicine, Argentina; Department of Psychiatry and Behavioral Sciences, Naresh K. Vashisht College of Medicine, Texas A&M University, College Station, Texas; Laureate Institute for Brain Research, Tulsa, Oklahoma; Grupo de Investigación en Neurociencias Aplicadas a las Alteraciones de la Conducta (Grupo INAAC), Instituto de Neurociencias Fleni-CONICET (INEU), Argentina; Departamento de Física, Facultad de Ciencias Exactas y Naturales, Universidad de Buenos Aires, Argentina

**Author notes:** **Correspondence:** Salvador M. Guinjoan, M.D., Ph.D., Division of Neuropsychiatry and Geropsychiatry, Department of Psychiatry and Behavioral Sciences, Naresh Vashisht College of Medicine, Texas A&M University. Shared authorship (Mirta F. Villarreal, Salvador M. Guinjoan).

## Abstract

**Background:** Depression is associated with risk for late-onset Alzheimer’s disease (LOAD), but its underlying pathogenesis in at-risk individuals remains unclear. We examined multimodal imaging correlates of depressive symptoms in cognitively normal middle-aged offspring of patients with LOAD (O-LOAD) compared with control individuals without LOAD history up to a 4^th^ degree of kinship (HC).

**Methods:** Participants (n=58; 52±3 years; 74% female) underwent assessment with the Beck Depression Inventory-II (BDI), structural MRI, resting-state fMRI, FDG-PET, and PiB-PET. Resting-state fMRI data were available for 28 O-LOAD and 24 HC; PET data for 24 O-LOAD and 22 HC. General linear models tested associations between imaging measures and BDI, including group interactions.

**Results:** In O-LOAD, higher BDI scores were associated with reduced cortical thickness in the left postcentral gyrus. Resting-state fMRI revealed significant group-by-BDI interactions involving cingulate and orbitofrontal networks. In O-LOAD, greater depressive symptom severity was associated with reduced cingulate connectivity across distributed corticolimbic, prefrontal, insular, occipital, and cerebellar regions (β range −0.10 to −0.18). In HC, depressive symptoms were associated with reduced right orbitofrontal and somatosensory–medial orbitofrontal connectivity (β=−0.13), with divergent patterns of cingulate connectivity. FDG-PET showed no significant associations with depressive symptoms. PiB-PET demonstrated regionally specific associations between amyloid signal and BDI in HC, involving an inverse pattern in anterior and posterior insular cortices.

**Conclusions:** Depressive symptoms in middle-aged individuals at familial risk for LOAD are associated with distinct structural and functional alterations, involving circuitry subserving salience and reward, and suggesting early network-level mechanisms linking affective symptoms with vulnerability to neurodegeneration.

## Introduction

Depression and neurocognitive disorders exhibit a complex epidemiological and phenomenological interaction (1). Burgeoning evidence indicates that depression increases the risk of both all-cause dementia and late-onset Alzheimer’s disease (LOAD) in a dose-dependent fashion, particularly when depression appears in middle age (2–6), while depressive symptoms are prevalent among patients with prodromal and established LOAD (7). The etiological basis for this interaction appears to be multifactorial, and includes shared vulnerability (including genetic factors), bidirectional causation, and possibly overlapping neurobiological disease pathways (8–10). Older persons newly diagnosed with major depressive disorder (MDD) are at maximum risk of receiving a clinical diagnosis of dementia within six months, a risk that remains elevated for up to two decades after the initial depression diagnosis (11). In turn, depression has been recognized as a modifiable risk factor for dementia in midlife, accounting for at least 3% of the population-attributable risk (12). A lifetime history of MDD, but particularly late-onset or recurring depression, significantly increases the risk of LOAD (1,13). The global prevalence of MDD among older adults was 13.3% (7); while reported prevalence of MDD is lower in older compared to younger adults (14), MDD among older adults is often underdiagnosed and undertreated, possibly reflecting stigma and atypical clinical presentations.

From a pragmatic standpoint the identification of mechanisms underlying symptoms of depression that herald progression to a chronic neurodegenerative condition like LOAD could have a critical impact in the field of mental health and neurocognitive disorders of older age. In this regard, preclinical LOAD, defined by biomarker abnormalities, may precede clinical diagnosis of dementia by 15-20 years (15). Currently, β-Amyloid and Tau biomarkers are typically assessed through advanced brain imaging techniques including positron emission tomography (PET) and magnetic resonance imaging (MRI), and the same applies to measurement of amyloid beta isoforms (Aβ) and phosphorylated tau in cerebrospinal fluid (CSF). While these imaging and CSF-based variables are reliable biomarkers for LOAD diagnosis and staging, their cost, invasiveness, and limited accessibility constrain their widespread and routine implementation (15). Further, despite recent significant advances (16), reliability and validity of blood-based biomarkers, which are minimally invasive, remain under investigation (17). These limitations underscore the need for noninvasive behavioral and neurobiological markers that may signal early pathophysiological processes resulting in LOAD.

In this context, persons at risk for LOAD have been extensively studied in older age (i.e., over 65 years of age) while still asymptomatic (18–24), but there is limited information on mechanisms of depressive symptom formation in cognitively normal persons with a family history of LOAD in middle age, i.e., more than a decade prior to the expected onset of dementia symptoms. In particular, there is a paucity of studies phenotyping such individuals with the help of multimodal neuroimaging, alongside appropriately matched healthy controls without a family history of LOAD. The inclusion of such comparison group would allow the identification of early, distinct depressive symptom mechanisms in persons who are at risk of LOAD by virtue of having at least one affected parent (a significant risk factor for LOAD, second only to advancing age, 25-27). In the present study we sought to identify multimodal neuroimaging correlates of depressive symptomatology in cognitively normal offspring of patients with probable late-onset Alzheimer’s disease (O-LOAD), and compare them with a group of demographically similar individuals but without evidence of LOAD in family members up to a 4^th^ degree of kinship (HC). Specifically, we sought to phenotype this group of individuals with structural (cortical thickness) and functional (resting-state connectivity) magnetic resonance imaging (MRI) variables, and with both fluorine-18 deoxyglucose (FDG-PET) and Pittsburgh compound B (PiB-PET) positron emission tomography, to observe regional abnormalities in brain metabolism and in vivo β-Amyloid deposition, respectively. We hypothesized that mechanisms of depressive symptoms are distinct in middle aged individuals who are genetically predisposed to LOAD, and reasoned that the current diagnostic approach to depression is insensitive to neurodegenerative behavioral phenocopies. We thus predicted that depression symptoms in these groups would be associated with (1) group-specific cortical thinning, (2) reduced resting-state fMRI connectivity within corticolimbic regions previously shown by our group and others to be structurally compromised in O-LOAD (28), (3) reduced regional cerebral metabolism as assessed by FDG-PET, particularly in mesial temporal and cuneus regions, or (4) increased insoluble β-amyloid deposition as measured by PiB-PET, in regions germane to depression pathophysiology.

## Materials and Methods

### Study Sample and Design

We report on a sample of offspring of LOAD (O-LOAD) and comparable individuals with no family history of LOAD up to a 4^th^ degree of kinship which has been described in detail elsewhere (28,29). The sample was composed of 28 O-LOAD and 24 HC participants for which fMRI data were available (Table 1). In this group, PiB-PET and FDG-PET images were available for 24 O-LOAD and 22 HC participants. All assessments were obtained within three months of recruitment. Subjects received no financial compensation for their participation in this study. The study protocol was approved by the Bioethics Committee of the FLENI Foundation and conducted in accordance with the Declaration of Helsinki. All participants provided written informed consent prior to participation.

**Table 1.** Demographic and clinical characteristics.

|  | HC | O-LOAD | Test statistic | p-value |
| --- | --- | --- | --- | --- |
|  | (N=26) | (N=32) |  |  |
| Female | 21 (80.8%) | 22 (68.8%) | $\chi^2 = 1.08$ | 0.299 |
| Age (yrs) | 50.3 (8.3) | 54.3 (7.6) | t = -1.92 | 0.060 |
| Education (yrs) | 17.8 (2.9) | 17.1 (3.1) | t = 0.87 | 0.390 |
| Hachinski Ischemic Score | 0.8 (1.0) | 1.2 (1.3) | t = -1.24 | 0.222 |
| BDI II | 8.1 (7.4) | 8.8 (6.4) | t = -0.36 | 0.717 |
| Estimated IQ | 107.5 (6.2) | 105.1 (6.5) | t = 1.41 | 0.165 |
| RAVLT Learning Curve | 48.3 (9.1) | 44.0 (7.4) | t = 1.93 | 0.060 |
| RAVLT Delayed Recall | 10.5 (2.1) | 8.6 (3.1) | t = 2.66 | **0.010** |
| RAVLT Recognition | 14.0 (1.1) | 13.0 (1.9) | t = 2.31 | **0.025** |
| Psychological treatment |  |  |  |  |
| No | 2 (7.7%) | 13 (40.6%) | Fisher's exact | **0.004** |
| Current | 8 (30.8%) | 2 (6.3%) |  |  |
| Past | 13 (50.0%) | 16 (50.0%) |  |  |
| Psychiatric treatment |  |  |  |  |
| No | 16 (61.5%) | 24 (75.0%) | Fisher's exact | 0.517 |
| Current | 4 (15.4%) | 2 (6.3%) |  |  |
| Past | 3 (11.5%) | 4 (12.5%) |  |  |
| Dyslipidemia | 5 (19.2%) | 12 (37.5%) | $\chi^2 = 2.25$ | 0.134 |
| Diabetes | 0 (0%) | 0 (0%) | Fisher's exact | 1.000 |
| Hypthiroidism | 6 (23.1%) | 4 (12.5%) | Fisher's exact | 0.307 |
| Cardiovascular | 3 (11.5%) | 4 (12.5%) | Fisher's exact | 1.000 |
| Neurologic | 3 (11.5%) | 3 (9.4%) | Fisher's exact | 1.000 |
| Autoimmune disorders | 1 (3.8%) | 1 (3.1%) | Fisher's exact | 1.000 |
| Other disorders | 0 (0%) | 6 (18.8%) | Fisher's exact | **0.036** |
| Lifetime smoking | 9 (34.6%) | 9 (28.1%) | $\chi^2 = 0.37$ | 0.542 |
| Antihypertensives | 0 (0%) | 5 (15.6%) | Fisher's exact | 0.059 |
| Benzodiazepines | 6 (23.1%) | 4 (12.5%) | Fisher's exact | 0.307 |
| SSRI | 3 (11.5%) | 3 (9.4%) | Fisher's exact | 1.000 |
| Anticonvulsants | 0 (0%) | 3 (9.4%) | Fisher's exact | 0.245 |
| Other medications | 7 (26.9%) | 12 (37.5%) | $\chi^2 = 0.66$ | 0.416 |

Values are presented as mean (SD) or n (%). P-values were calculated using Student's t-test for continuous variables and chi-square or Fisher's exact test for categorical variables.

Inclusion criteria for the O-LOAD group were: (a) having at least one parent diagnosed with probable late-onset Alzheimer’s disease according to DSM-5 criteria; (b) age between 40 and 65 years at recruitment; (c) more than seven years of formal education; and (d) self-reported absence of clinical depression (some participants were determined to suffer from major depression by a consulting psychiatrist (SMG), as reported in Table 1). Although postmortem confirmation was not available for affected parents, participants provided detailed clinical information including medical diagnosis in the medical records for their family members. For parents who had not been treated at the FLENI Foundation (n = 5), the diagnosis of LOAD was confirmed by an independent clinician. In addition to clinical diagnosis, structural MRI scans were available for the parents of 15 participants, allowing confirmation of atrophic patterns consistent with LOAD and the absence of significant cerebrovascular pathology. Among these cases, three parents also showed positive amyloid deposition on PiB-PET imaging. None of the participants had both parents diagnosed with LOAD.

The HC group met the same inclusion criteria, with the exception that they reported no history of Alzheimer’s disease or other neurodegenerative disorders up to a fourth degree of kinship.

Exclusion criteria for all participants included: (a) compromised intellectual ability measured by a complete neuropsychological battery that evaluated all cognitive domains (estimated IQ, attention, memory, language, executive functions); (b) evidence of current progressive neurologic disease or medical conditions likely to impair cognitive function (c) history of substance abuse (including alcohol, marijuana, stimulants, benzodiazepines, cocaine, or other illicit drugs); and (d) a Hachinski score > 7, to screen out individuals with potentially significant cerebrovascular compromise (30). Figure 1 shows an overall description of the study design.

**Figure 1:**
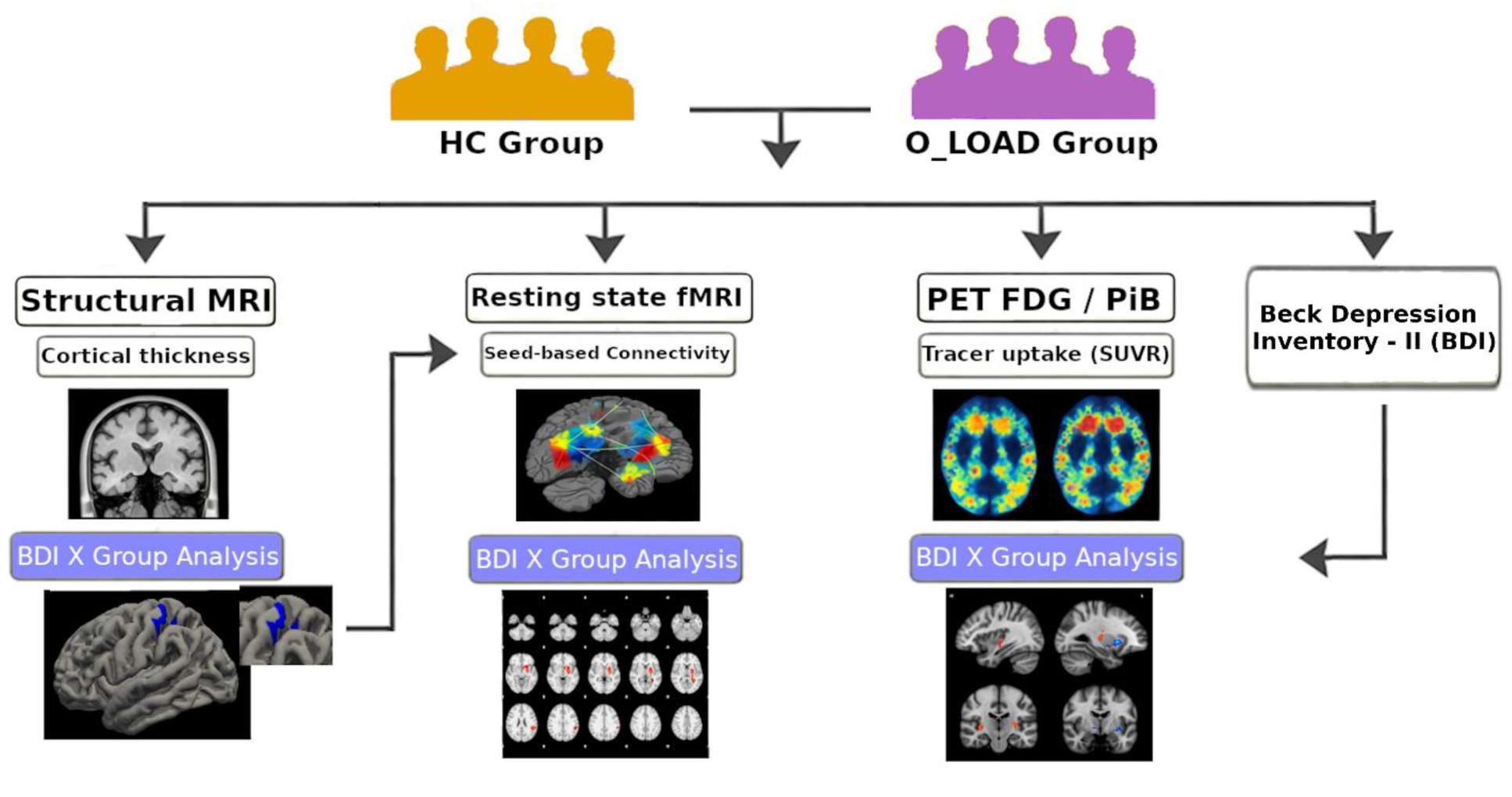
Workflow of multimodal neuroimaging analyses, including cortical thickness from structural MRI, seed-based functional connectivity from resting-state fMRI, and PET measures, in relation to BDI × group effects. O-LOAD: offspring of late-onset Alzheimer’s disease patients; HC: controls without a family history of late-onset Alzheimer’s disease up to a 4^th^ degree of kinship; BDI: Beck Depression Inventory-II.

### Cognitive assessment

Potential participants were first screened using the WAT-BA to obtain an estimated IQ score (31). Individuals who met the inclusion criteria completed a comprehensive neuropsychological assessment to confirm the absence of cognitive impairment and to ensure intellectual functioning within the normal range. This battery consisted of standardized neuropsychological tests that are well validated and widely used in clinical practice, assessing attention, memory, language, and executive function cognitive domains; therefore, detailed descriptions are not provided here. The specific tests administered have been reported in detail elsewhere (32).

Depressive symptoms were assessed using the self-administered Beck Depression Inventory scale–II (BDI; 33). Caregiver burden was evaluated using the 22-item Zarit Burden Interview (34). Among O-LOAD participants, two individuals reported being active caregivers at the time of recruitment; the remaining O-LOAD parents were either institutionalized or deceased. Of those two caregivers, only one showed scores indicative of significant caregiver burden, based on a cutoff score of 48 in the Zarit Burden Interview (35). None of the control participants reported caregiving responsibilities.

All neuropsychological assessments were administered by a trained neuropsychologist (C.A.) who was blind to the participants’ group status (O-LOAD vs. HC). Performance on all neuropsychological measures was within normal limits according to local normative data. Accordingly, none of the participants met diagnostic criteria for minor or major neurocognitive disorder.

### MRI acquisition

MRI images were obtained with a 3T General Electric HDxt MRI scanner with an 8-channel head coil. A structural T1-weighted MRI was obtained with the 3D fast SPGR-IR sequence (166 contiguous slices at TR/TE = 7.3/3.0 ms, flip angle 8°, FOV/slice thickness 260/1.2 mm, image matrix 256 × 256). Resting state functional imaging (rs-fMRI) signals were acquired with the EPI-GRE sequence (33 contiguous slices in the AC–PC plane, TR/TE=2000/30 ms, flip angle 90°, FOV=24 cm, voxel volume = 3.75 × 3.75 × 4 mm^3^, acquisition matrix = 64 x 64) along 200 volumes per subject.

### [^18F] FDG synthesis and PET/CT acquisition

PET data were acquired on a GE Discovery 690 PET/CT scanner (GE Healthcare). [^18F] FDG was synthesized on-site using a FASTlab automated synthesizer (GE Healthcare) with single-use disposable cassettes. [^18F] Fluoride was produced by proton irradiation of enriched [^18O] water (2.5–5 mL) in a cyclotron. Typical production yields ranged from 1500 to 7000 mCi. Total synthesis time was approximately 25 min, yielding ∼15 mL of injectable solution after standard quality control.

Participants fasted for ≥6 h and had blood glucose levels <130 mg/dL prior to injection. Tracer uptake occurred in a quiet, dimly lit room. PET acquisition began 30–40 min after intravenous [^18F] FDG administration. Dynamic 3D PET data were acquired over approximately 30 min.

### [^11C] PiB synthesis and PET/CT acquisition

[^11C] PiB was synthesized using a GE TRACERlab FX C Pro automated module from [^11C]CO₂. Radiochemical purification was performed by reverse-phase HPLC using 0.009 M sodium citrate in ethanol/water (60/40, v/v) as the mobile phase at a flow rate of 3 mL/min. Elution was monitored with UV and radioactivity detectors in series.

The [^11C] PiB fraction (retention time 11–12 min) was collected by automated valve switching, diluted with 30 mL of saline to reduce ethanol content, and sterile-filtered, resulting in a final volume of 36 mL. A dose of 10 mCi (weight-adjusted) was administered intravenously. PET acquisition commenced 50 min post-injection, with dynamic 3D imaging performed over 20 min.

### MRI T1 and PET image preprocessing

Cortical reconstruction was performed using FreeSurfer image analysis software (version 6.0; http://surfer.nmr.mgh.harvard.edu/). The standard FreeSurfer processing pipeline includes removal of non-brain tissue (36), Talairach transformation, and automated segmentation of subcortical structures (37,38). Additional steps comprise intensity normalization (39), topological correction (40,41), and surface deformation driven by intensity gradients to accurately delineate the gray matter/white matter (GM/WM) and gray matter/cerebrospinal fluid boundaries, corresponding to regions of maximal intensity change at tissue class transitions (42–44). Preprocessing was performed independently for each hemisphere. All reconstructed surfaces were visually inspected and manually corrected to ensure anatomical accuracy. Surface maps were subsequently smoothed prior to statistical analysis.

Whole-brain cortical thickness was quantified at each vertex as the shortest distance between the GM/WM boundary and the pial surface. Thickness estimation relied on both absolute signal intensity and spatial intensity gradients across tissue classes.For inter-subject alignment and group-level analyses, individual cortical thickness maps were registered to a common spherical atlas (45), enabling the generation of surface-based measures including cortical thickness, curvature, and surface area.

PET images were processed in conjunction with volumetric T1-weighted MRI data, and analyzed using PETSurfer scripts obtaining the corresponding PET-PiB and PET-FDG intensity maps. Positron emission tomography (PET) images pipeline using Petsurfer script included: (a) generation of high-resolution segmentations used to run partial volume correction (PVC, 46), (b) co-registration of PET images to individual T1-weighted MRI scans, (c) application of established PVC methods (47–50), and (d) projection of corrected cortical and subcortical uptake values onto individual subject surfaces. The pons was selected as the reference region due to low susceptibility to partial volume effects, relative resistance to atrophy in Alzheimer’s disease, and common use in FDG and amyloid PET studies.

### Resting-state fMRI image processing

#### Preprocessing

rs-fMRI data were analyzed using the FMRIB Software Library (FSL, version 6.0.6.2; FMRIB Analysis Group, University of Oxford). Preprocessing was performed using FEAT (FMRIB’s Expert Analysis Tool) and included brain extraction using BET, motion correction with MCFLIRT, and spatial normalization to MNI standard space using linear (FLIRT) and nonlinear (FNIRT) registration. Temporal high-pass filtering was applied with a cutoff of 100s, and spatial smoothing was performed using a Gaussian kernel with an 8 mm full width at half maximum.

#### Seed-based analysis

Seed-based analysis was performed to assess resting-state functional connectivity (51). Four regions of interest (ROIs) were selected as seeds, based on their trophic compromise in O-LOAD (28), namely bilateral cingulate (CC) and orbitofrontal (OFC) cortices. These ROIs were obtained from the Harvard Oxford atlas within FSL software. An additional ROI was included as a seed, based on significant cortical thickness correlates of depressive symptoms identified in the structural MRI analysis in the present study (left primary somatosensory cortex). For each seed, the mean BOLD time series was extracted from individual rs-fMRI data and correlated with the time series of all other brain voxels, generating whole-brain functional connectivity maps.

### Statistical analysis

#### Cortical statistical analysis

To assess the association between BDI and cortical thickness, a surface-based group analysis was performed using FreeSurfer’s general linear model framework. Cortical thickness maps were entered as dependent variables with BDI included as a continuous covariate of interest. The design matrix further included the group factor and the BDI × Group interaction term, allowing the examination of group-specific associations between BDI and cortical thickness.

Correction for multiple comparisons was conducted using Monte Carlo simulation with a cluster-wise significance threshold of p < 0.05 and a vertex-wise (cluster-forming) threshold of p < 0.05. Significant clusters were visualized on semi-inflated cortical surface maps, with warm colors indicating positive associations and cool colors indicating inverse associations between cortical thickness and BDI scores.

#### Seed-based statistical analysis

To examine the effect of BDI on functional connectivity between the selected ROIs and the whole brain, group-level analyses were performed using a general linear model (GLM) implemented in FSL (52). Individual whole-brain connectivity maps derived from each seed were entered as dependent variables, with BDI included as a continuous covariate of interest. The design matrix further included the group factor and the BDI × Group interaction term, allowing the examination of group-specific associations between BDI and functional connectivity. Correction for multiple comparisons was performed using cluster extent–based thresholding (53). Statistical Z images were thresholded at Z > 2.3, with a cluster-level significance threshold of p = 0.05 (54).

For visualization and characterization of the interaction effects, mean connectivity values were extracted from significant clusters for each participant and entered into linear regression models to estimate simple slopes within each group. Results were visualized using scatter plots with fitted regression lines. Regressions and plots were performed with R software (version 4.4.1; R Core Team, Vienna, Austria).

#### PET statistical analysis

To investigate the relationship between PET measures and BDI scores, we employed the same GLM framework described above. Regional cortical surface–based and subcortical volumetric PET uptake values (^11^C-PiB or ^18^F-FDG) were entered as dependent variables, with BDI included as a continuous covariate of interest alongside the group factor and their interaction term. Correction for multiple comparisons was conducted using Gaussian random field (GRF) theory, with a cluster-wise significance threshold of p < 0.05 and a vertex-wise (cluster-forming) threshold of p < 0.001.

## Results

### Participant Characteristics

Both groups were comparable regarding sex, age, baseline cognitive functioning, and years of education (Table 1). BDI scores were on average within a non-clinically significant range in both groups (Table 1), but three controls and seven O-LOAD participants reported a BDI>12.

### Structural and Functional Magnetic Resonance Imaging

A significant negative association between cortical thickness and BDI scores was observed in a discrete area of the left postcentral gyrus within the O-LOAD group only (Figure 2), overlapping with the primary somatosensory cortex.

**Figure 2:**
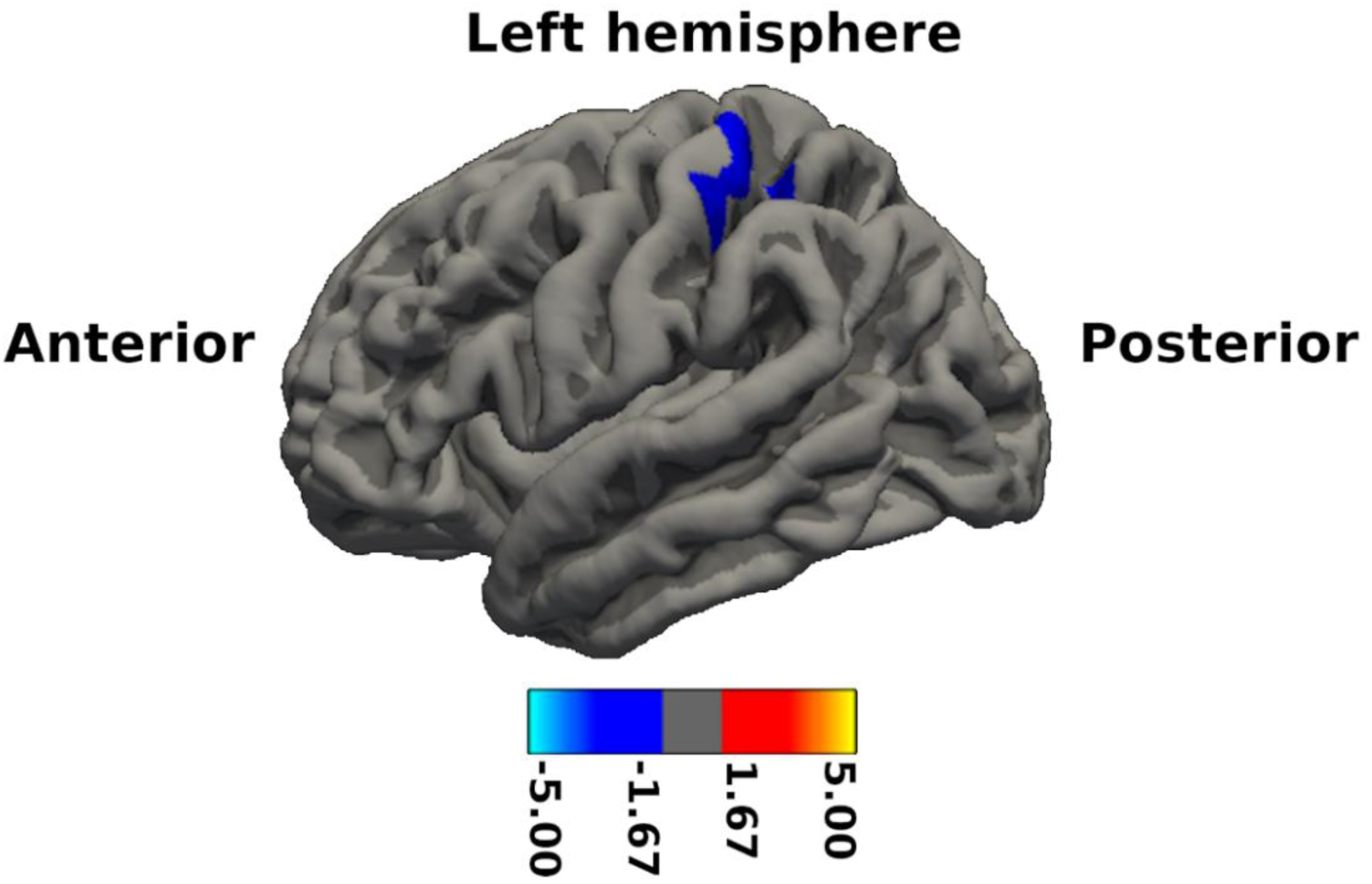
Association between BDI scores and cortical thickness in O-LOAD participants. Blue colors indicate a negative association between BDI scores and cortical thickness in a cluster located in the left postcentral gyrus. The statistical map is displayed as −log(p) values and thresholded at p < 0.05 using cluster-wise Monte Carlo correction. BDI: Beck Depression Inventory-II; O-LOAD: offspring of late-onset Alzheimer’s disease.

Regarding resting-state functional MRI connectivity, significant BDI × Group interaction effects were observed for three seed regions previously reported to show cortical thinning in O-LOAD (28). Post-hoc extraction of mean connectivity values from significant clusters revealed differential associations with BDI across groups, with opposite slopes in all ROIs (Figures 3–5; Table 2).

**Figure 3:**
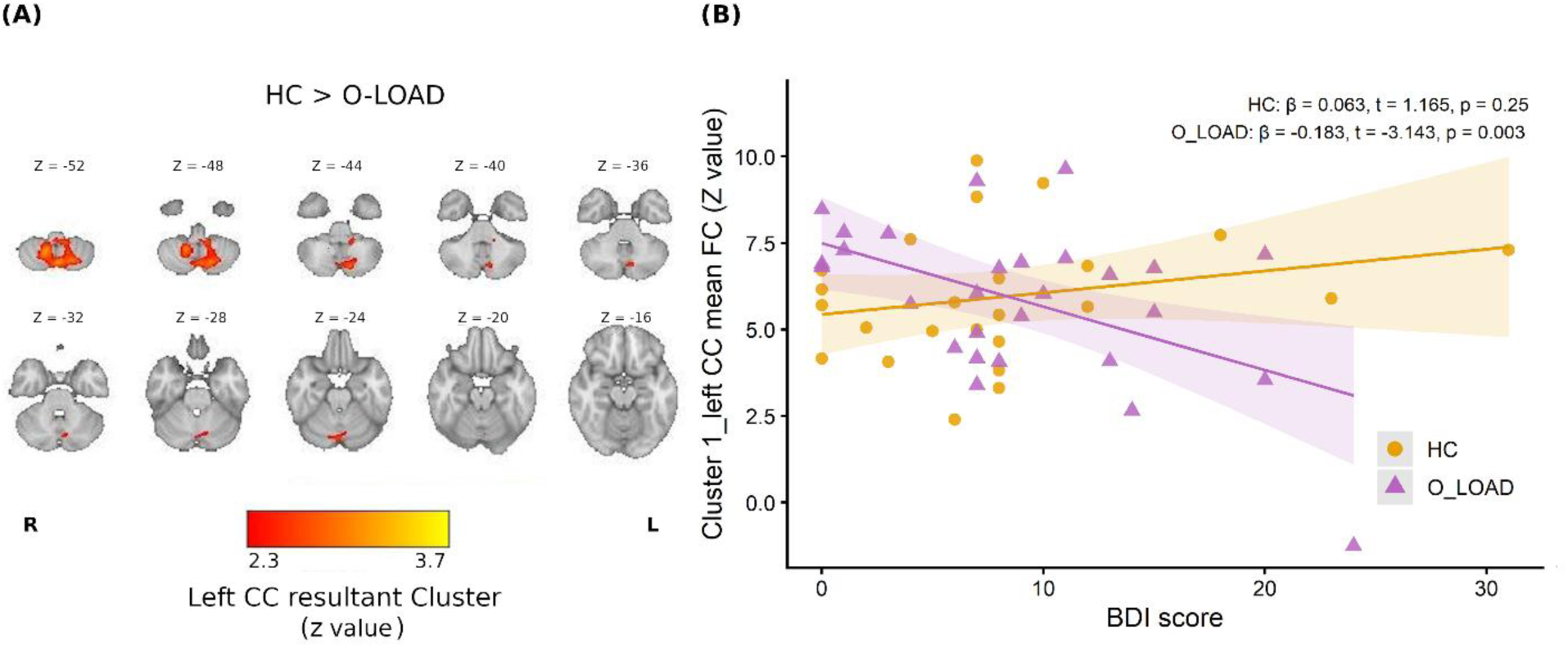
Association between BDI scores and functional connectivity of the left cingulate cortex. (A) Whole-brain maps showing the significant BDI × Group interaction (HC > O-LOAD) in seed-to-voxel connectivity with the left CC, (p_corrected < 0.05, z coordinates in MNI representation). (B) Scatter plots of BDI scores versus mean connectivity within the resultant cluster for O-LOAD and HC, showing a significant negative association for O-LOAD participants. BDI: Beck Depression Inventory; O-LOAD: offspring of late-onset Alzheimer’s disease; HC: controls without family history of Alzheimer’s disease up to a 4^th^ degree of kinship; FC: functional connectivity; CC: cingulate cortex.

**Figure 4:**
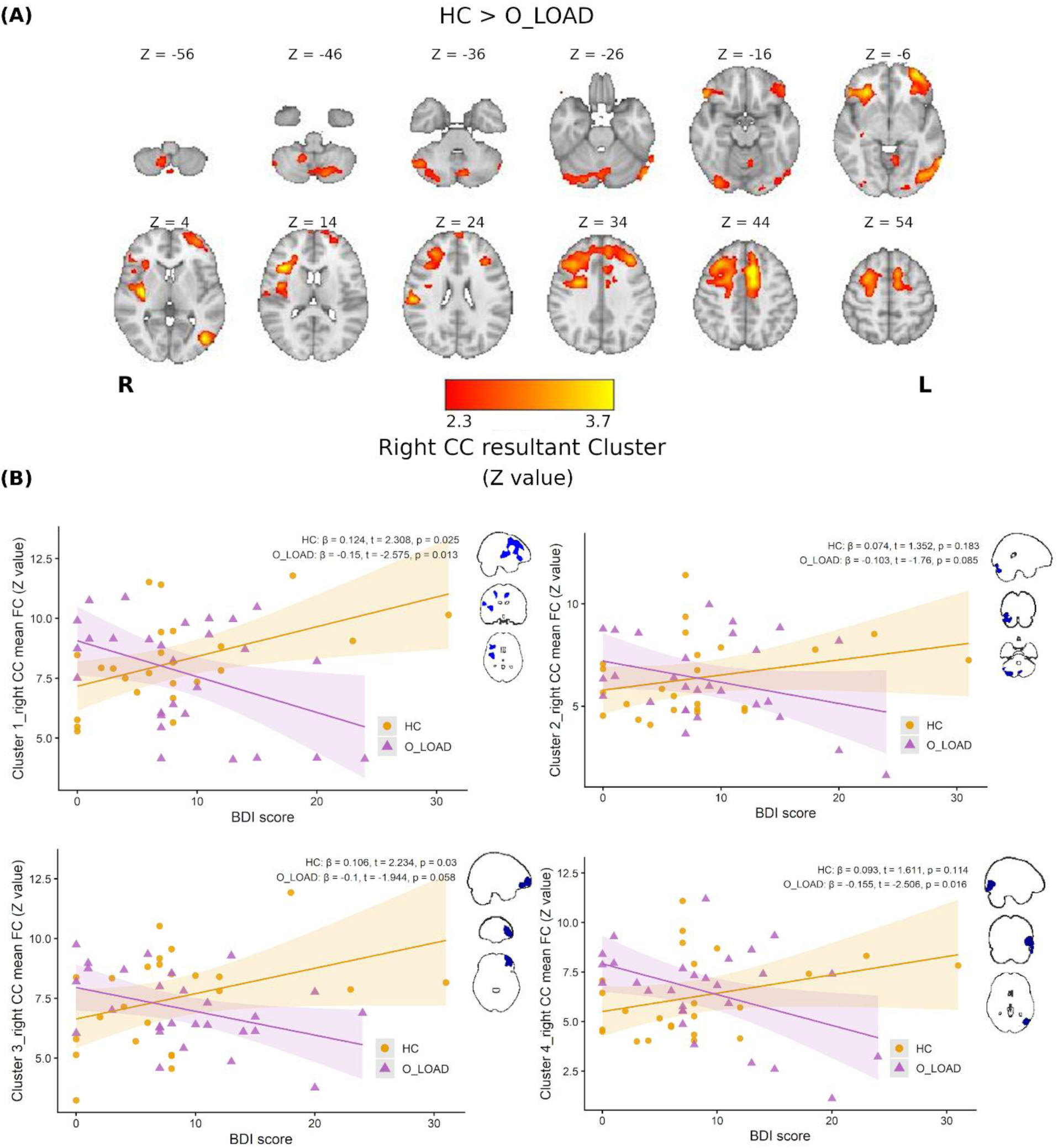
Association between BDI scores and functional connectivity of the right cingulate cortex. (A) Whole-brain maps showing the significant BDI × Group interaction (HC > O-LOAD) in seed-to-voxel connectivity with the right CC, (p_corrected < 0.05, z coordinates in MNI representation). (B) Scatter plots of BDI scores versus mean connectivity within the resultant clusters for O-LOAD and HC, showing a significant negative association for O-LOAD participants in clusters 1, 3, and 4, and a direct relationship for HC in clusters 1 and 3. BDI: Beck Depression Inventory; O-LOAD: offspring of late-onset Alzheimer’s disease; HC: controls without family history of Alzheimer’s disease up to a 4^th^ degree of kinship; FC: functional connectivity; CC: cingulate cortex.

**Figure 5:**
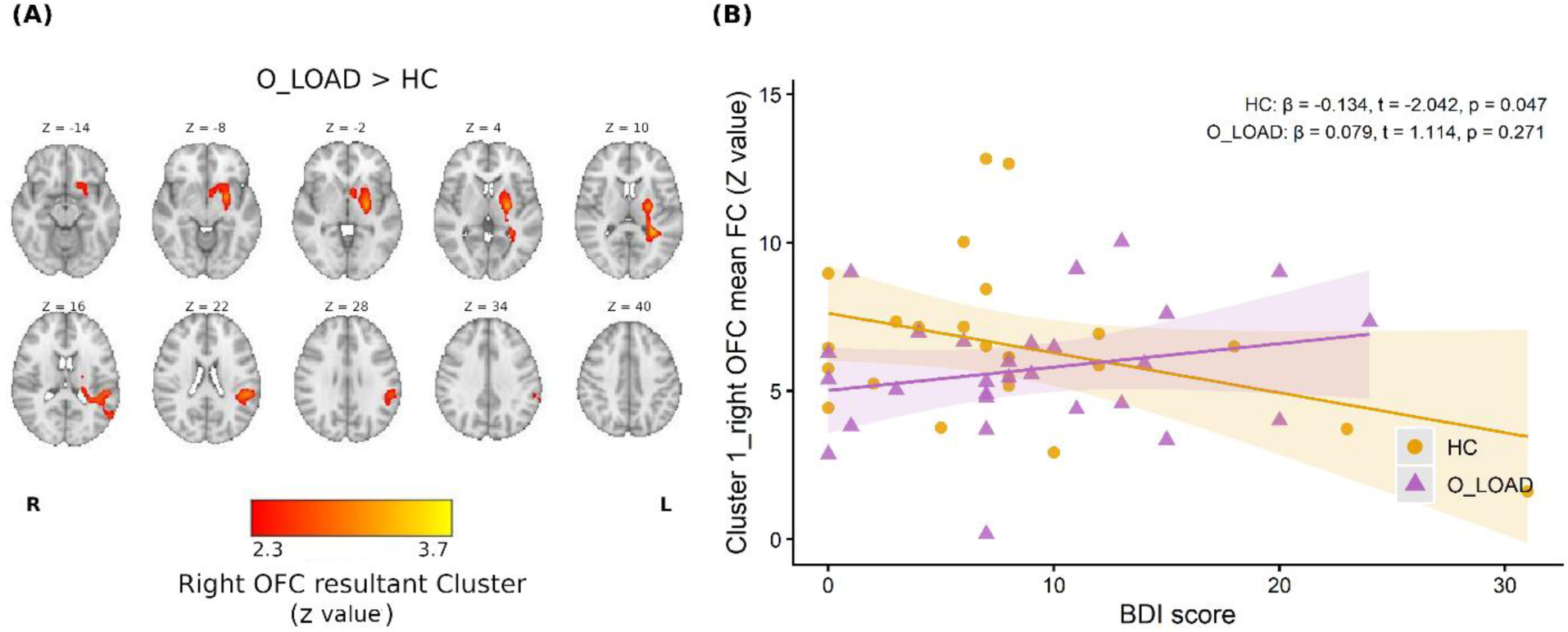
Association between BDI scores and functional connectivity of the right orbitofrontal cortex. (A) Whole-brain maps showing the significant BDI × Group interaction (O-LOAD > HC) in seed-to-voxel connectivity with the right OFC, (p_corrected < 0.05, z coordinates in MNI representation). (B) Scatter plots of BDI scores versus mean connectivity within the resultant cluster for O-LOAD and HC, showing a significant negative association for HC participants. BDI: Beck Depression Inventory; O-LOAD: offspring of late-onset Alzheimer’s disease; HC: controls without family history of Alzheimer’s disease up to a 4^th^ degree of kinship; FC: functional connectivity; OFC: orbitofrontal cortex.

**Table 2:**
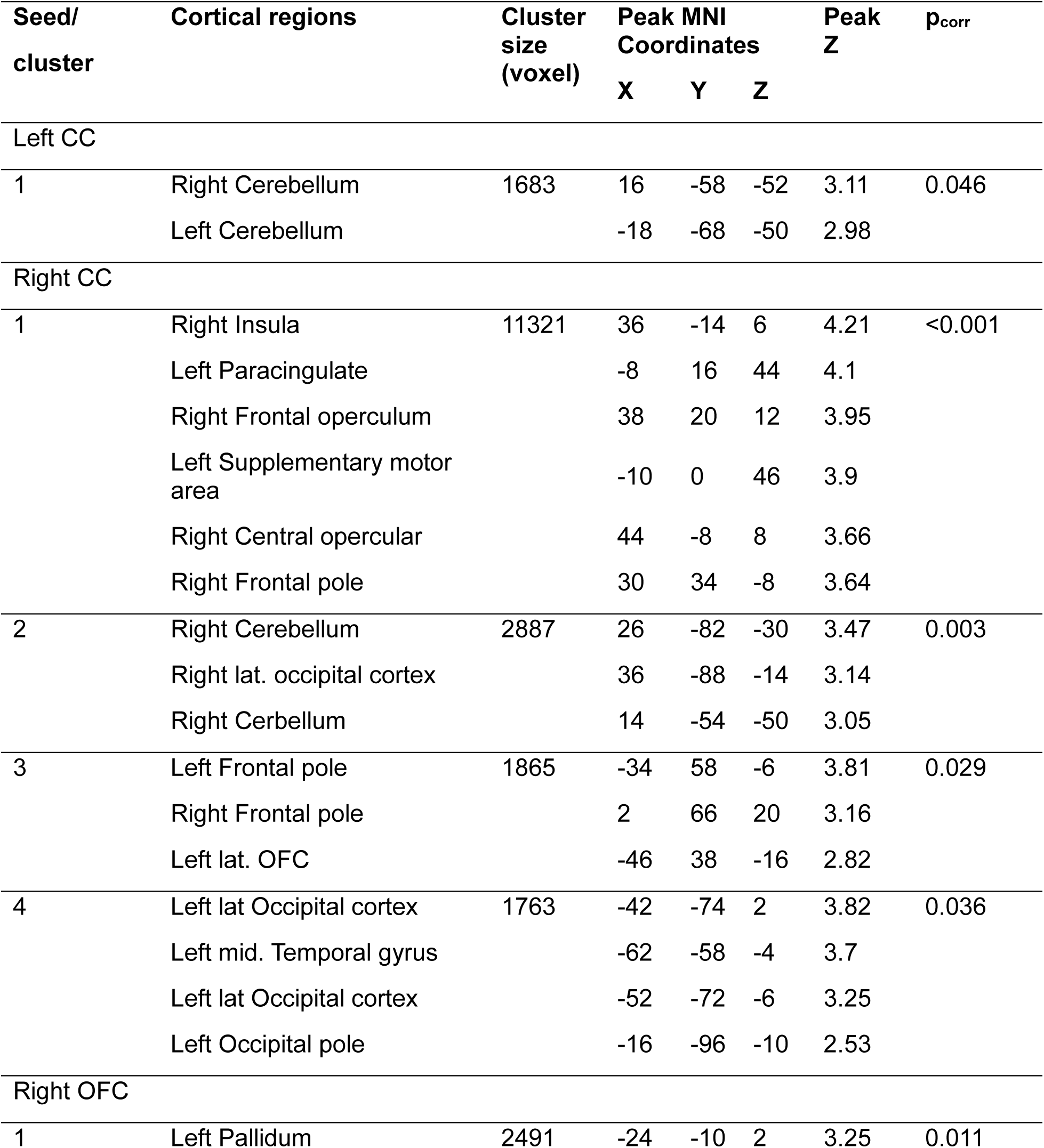
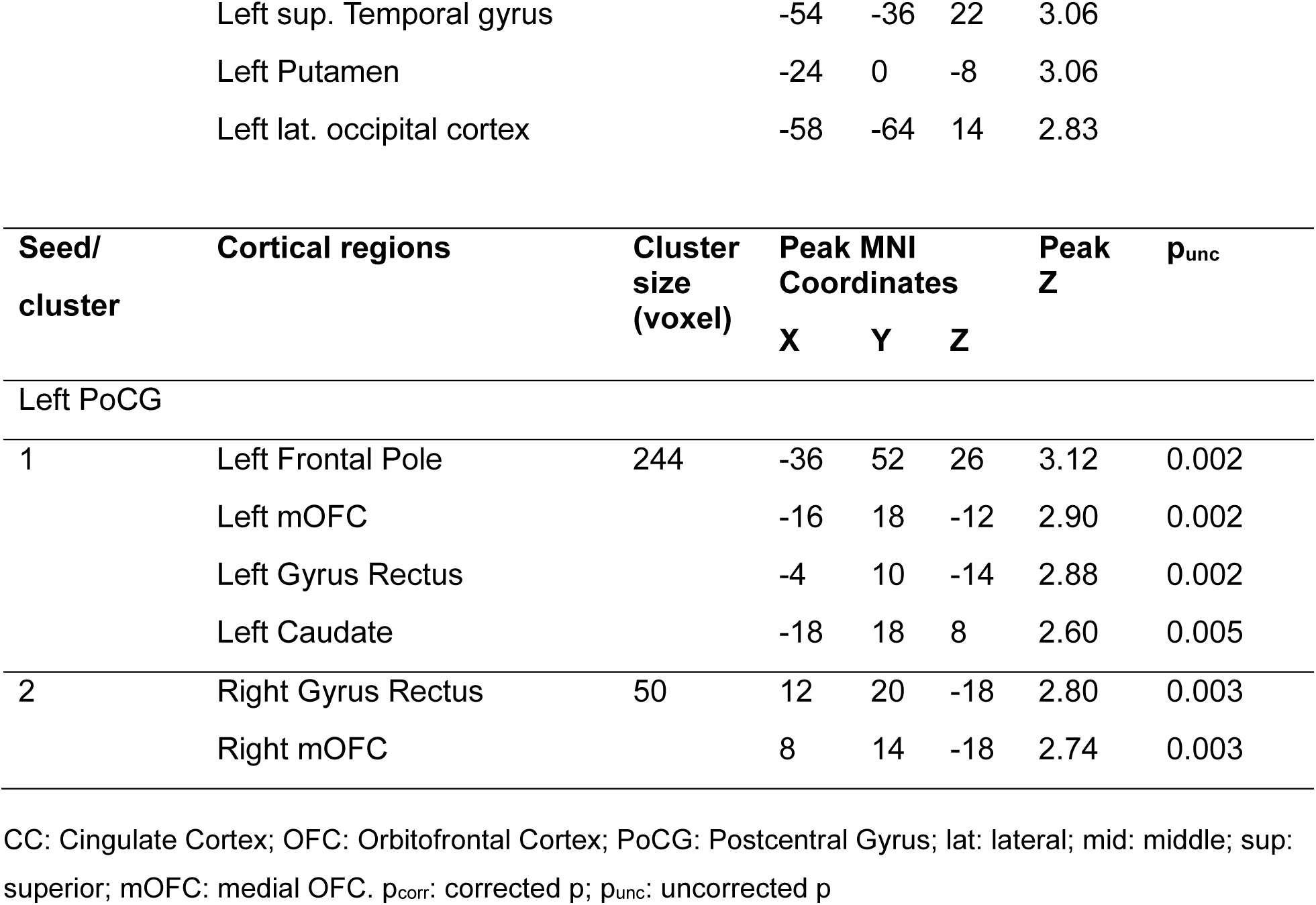
Clusters showing significant BDI × Group interactions in seed-based connectivity analyses.

For the left cingulum seed, whole-brain seed-to-voxel analysis revealed a significant BDI × Group interaction involving widespread archicerebellar and paleocerebellar regions, with a greater BDI-related slope in HC compared to O-LOAD. Post-hoc regression analyses showed a significant negative association between connectivity and depressive symptoms in O-LOAD, whereas no significant association was observed in HC participants (Figure 3, panels A and B; Table 2).

For the right cingulum seed, whole-brain analysis revealed four clusters showing a significant BDI × Group interaction (HC > O-LOAD; Figure 4, panel A; Table 2). These clusters were more extensive than those observed for the left cingulum and included bilateral dorsolateral prefrontal, lateral orbitofrontal, and right insular cortices (Cluster 1), right neocerebellar and occipital cortices (Cluster 2), left frontopolar cortex (Cluster 3), and left middle occipital region (Cluster 4). Post-hoc regression analyses demonstrated a significant negative association between functional connectivity and BDI scores in O-LOAD across all clusters. In HC participants, a significant positive association was observed in clusters 1, 3, and 4, whereas no significant association was found in cluster 2 (Figure 4, panel B).

Whole-brain analysis for the right OFC seed revealed a significant BDI × Group interaction (O-LOAD > HC) within a widespread left-hemispheric cluster encompassing the putamen, pallidum, nucleus accumbens, insula, inferior parietal lobule, supramarginal gyrus, and posterior temporal regions (Figure 5, panel A). Post-hoc regression analyses within this cluster showed a significant negative association between connectivity strength and BDI scores in HC participants, whereas no significant association was observed in O-LOAD (Figure 5, panels A and B; Table 2). Connectivity of the left OFC seed had no relationship with BDI in either group.

Based on the cortical thickness findings (Figure 2), a seed-to-voxel analysis was also conducted using the left postcentral region in relation to BDI in O-LOAD. This analysis revealed a BDI × Group interaction (O-LOAD > HC) in bilateral medial orbitofrontal cortex (mOFC) and gyri recti. This effect did not survive correction for multiple comparisons. Post-hoc regression analyses within the right mOFC/gyrus rectus cluster (Cluster 2) demonstrated a significant negative association between connectivity strength and BDI scores in HC participants, whereas no significant association was observed in O-LOAD (Figure 6, panels A and B; Table 2).

**Figure 6:**
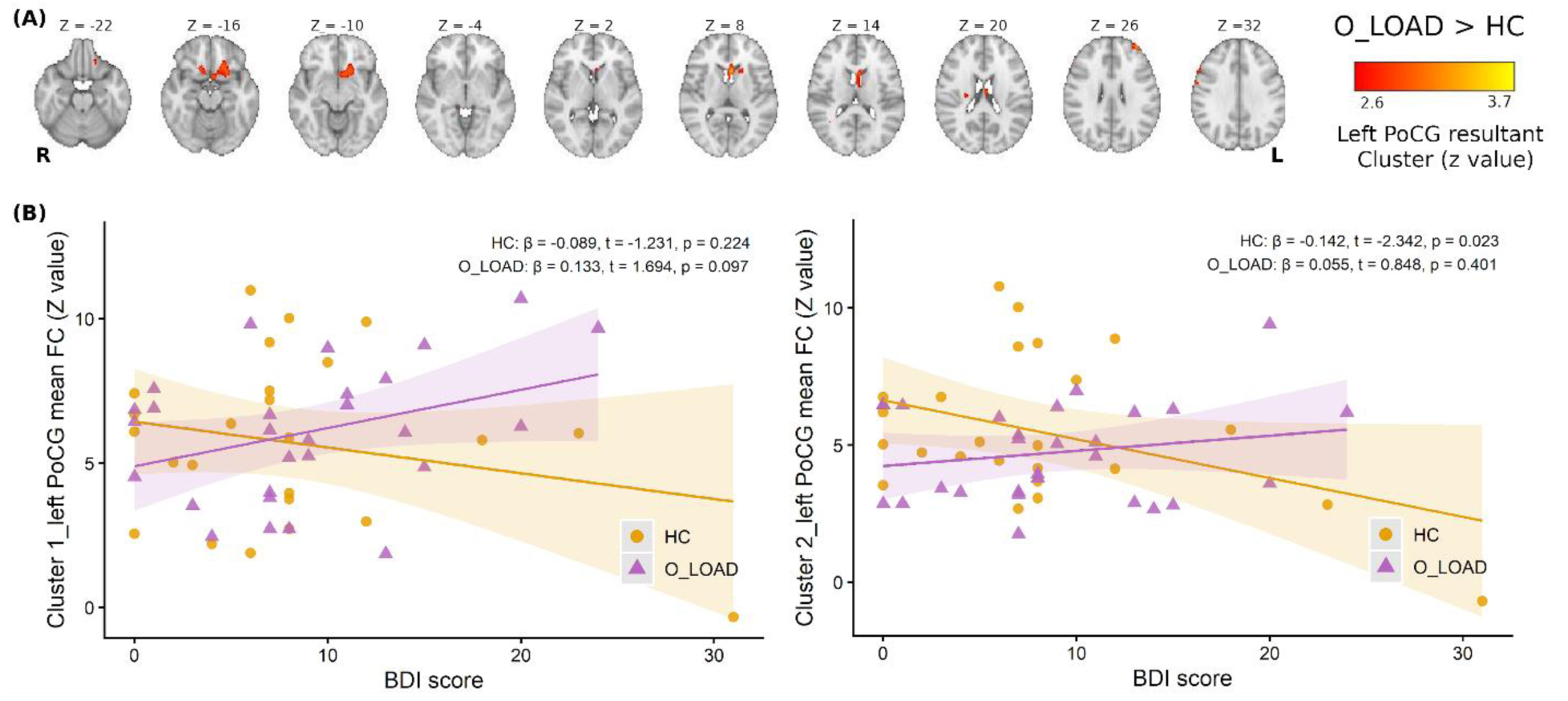
Association between BDI scores and functional connectivity of the left Postcentral gyrus. (A) Whole-brain maps showing the significant BDI × Group interaction (O-LOAD > HC) in seed-to-voxel connectivity with the right PoCG, (p_uncorrected < 0.005, z coordinates in MNI representation). (B) Scatter plots of BDI scores versus mean connectivity within the resultant clusters for O-LOAD and HC, showing a significant negative association for HC in the right medial orbitofrontal cortex (clusters 2). BDI: Beck Depression Inventory; O-LOAD: offspring of late-onset Alzheimer’s disease; HC: controls without family history of Alzheimer’s disease up to a 4^th^ degree of kinship; FC: functional connectivity; PoCG: postcentral gyrus.

### Positron Emission Tomography

PET-FDG analyses revealed no significant BDI × Group interactions or within-group associations with BDI scores. For PET-PiB, no significant BDI × Group interaction was observed after correction for multiple comparisons. However, exploratory analyses at an uncorrected threshold (p < 0.005) indicated that, within HC participants, PET-PiB uptake showed significant associations with BDI scores, displaying opposite-direction associations across anterior and posterior insular cortices. Posterior insular PET-PiB uptake was positively associated with BDI scores, whereas anterior insular regions exhibited an inverse association (Figure 7). Additionally, β-amyloid deposition in HC participants was inversely correlated with BDI scores in bilateral putamen and left posterior thalamus.

**Figure 7:**
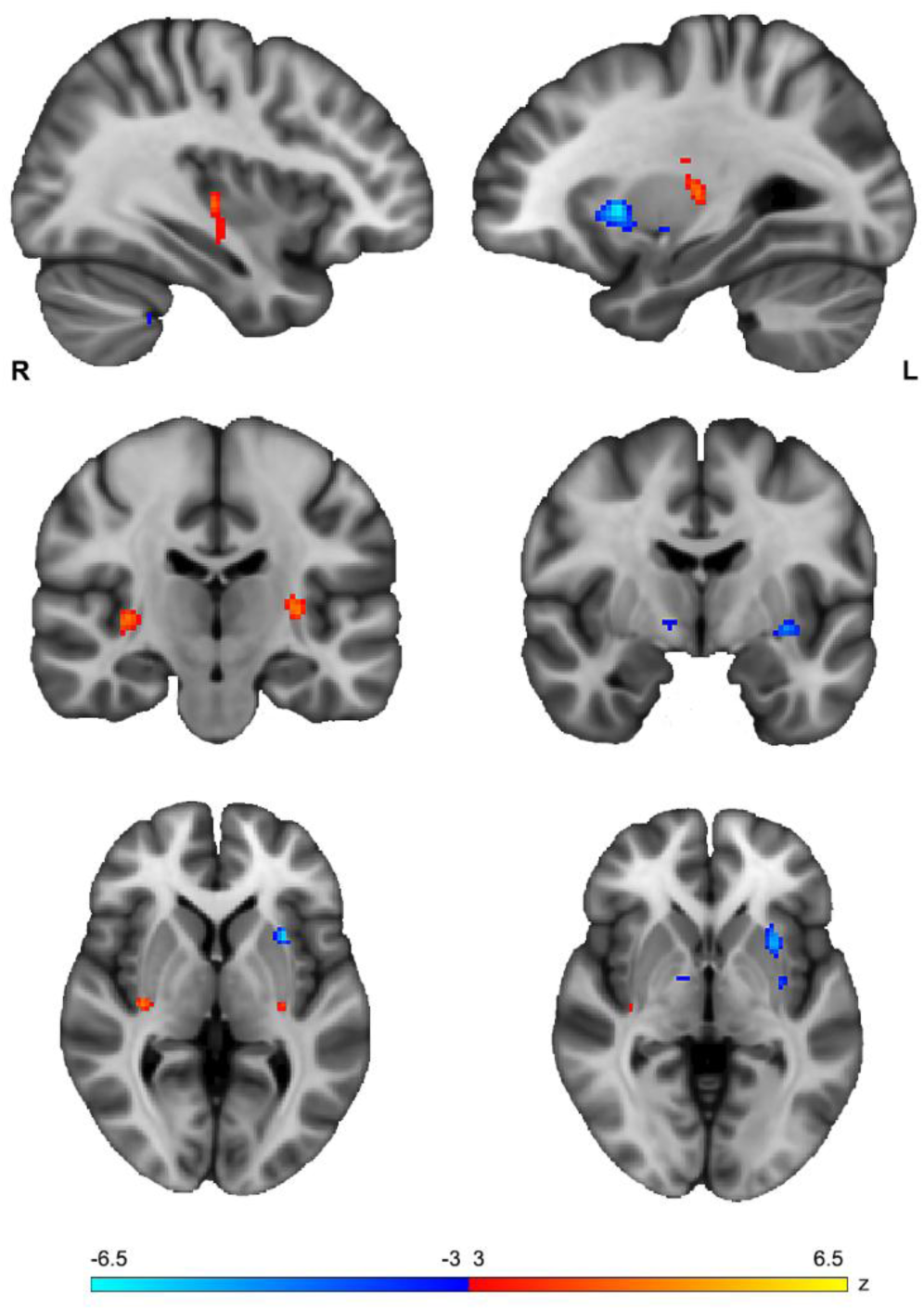
Association between BDI scores and brain β-amyloid deposition measured with PiB-PET in HC. Warm colors indicate positive associations between BDI scores and PiB uptake in the posterior right insula and left putamen. Cool colors indicate negative associations in the anterior left insula, left putamen, and right pallidum. MNI coordinates (x, y, z) of the slices shown correspond to (36, −19, 2) for the left column and (−28, −6, −2) for the right column. Statistical maps are displayed at an uncorrected threshold of p < 0.001. BDI: Beck Depression Inventory-II; HC: controls without family history of Alzheimer’s disease up to a 4^th^ degree of kinship.

## Discussion

Leveraging our prior observations on structural and functional imaging differences between cognitively normal, middle-aged adults with (O-LOAD) and without (HC) LOAD extended family history (28,29,55–58), in the present study we set out to explore the relationships between multimodal imaging variables and the presence of self-reported depressive symptomatology as assessed by BDI score in these groups. Our main findings are (1) an inverse relationship between BDI score and whole-brain connectivity in corticolimbic regions previously observed to be atrophic in O-LOAD compared with HC, (2) diminished left somatosensory cortex thickness in O-LOAD in relation to depressive symptoms, (3) a relationship between functional connectivity of this somatosensory cortex area with medial orbitofrontal cortices, and lower BDI scores, only in HC, (4) no significant relationships between regional metabolism as assessed with FDG-PET and BDI in either group, and (5) unexpectedly, a distinct, graded pattern of the relationship between PiB-PET signal in bilateral insular subregions, along with positive associations in bilateral putamina, and posterior left thalamus, which were present only among individuals with no family history of LOAD. To the extent of our knowledge, a distinct feature of our study is the comparison of O-LOAD, over a decade before the expected onset of any AD symptoms on average, with a matched sample of individuals who are also cognitively normal but without a close (4^th^ degree of kinship) family history of LOAD (HC). Findings point to both potential mechanisms of depression symptom development in the context of preclinical LOAD, and a relationship between β-Amyloid deposition and the self-reported presence of depressive symptoms in middle aged persons who do not have an increased family risk of LOAD.

Our results suggest a potential role of functional connectivity deficits of CC in O-LOAD (previously shown to be atrophic in comparison with HC) in the generation of self-reported symptoms of depression. Such connectivity deficits involved primarily the right cingulate cortex, and a series of regions germane to executive functioning, salience, visceral regulation, and higher order sensory processing (59,60). In general, faulty activation and connectivity of the CC has been consistently observed to be a feature of MDD (61,62). Consistent with these observations, O-LOAD participants in our study displayed an inverse relationship between connectivity of the right anterior cingulate cortex and depressive symptom severity assessed with BDI. The variety of cortical areas exhibiting this relationship probably reflects the nodal position of the anterior cingulate cortex (ACC) in the regulation of emotion, visceral function, executive function, assessment of rewarding clues, and related decision-making abilities that inform motivated behavior and pleasure (59–62). Regions thus disconnected from the ACC conceivably contribute to depressive symptoms in the specific group of middle-aged O-LOAD, suggesting that different etiological origins may converge on a similar pathophysiology of depression. In the case of our O-LOAD sample, we had previously observed significant cortical atrophy of the ACC in comparison with matched controls without a family history of LOAD, and now we observe that the functional correlate of such neurotrophic profile is poor connectivity akin to that observed in MDD, albeit with a different, possibly incipient neurodegenerative, origin. Along with other findings reviewed herein, these results suggest that depression in middle-aged individuals who are at familial risk of O-LOAD, merits further exploration as an early biomarker of neurodegeneration, possibly over a decade before the expected onset of symptoms. Glaubitz and coworkers have recently observed a similar dissociation but referred to in vivo Tau deposition in relation to BDI, yet in a sample with a wider age range, including much older individuals with positive LOAD biomarkers (63).

We observed an area of the postcentral gyrus in the dominant hemisphere, functionally overlapping with the primary somatosensory cortex, whose thickness was inversely related to depressive symptoms assessed with BDI in O-LOAD. Functional connectivity of this specific area with the OFC showed an inverse relationship with depressive symptom severity in persons without genetic predisposition to LOAD. The mOFC is well-characterized as a major hub in the canonical reward circuit affected in depression, and in particular, its connectivity to the somatosensory cortex may underlie the positive emotional coding of somatosensory input, which in turn may inform the output of psychomotor activity and behavior. If this is the case, faulty connectivity of somatosensory cortex and mOFC driven by incipient trophic phenomena in O-LOAD, may explain in part the presence of depressive symptomatology. Therefore, our findings in this region possibly warrant the exploration of hedonic dysfunction in late-onset depression to discriminate between primary depressive pathophysiology in the form of anhedonia, and symptoms originating from incipient neurodegeneration akin to apathy, possibly as an early biomarker of LOAD risk.

The absence of FDG-PET correlates of depressive symptoms in either study group was an unexpected finding which may obey two main reasons. First, regional activation differences may be subtle enough to not being detected at the group level in the relatively small sample that we have investigated. Second, any incipient neurodegenerative changes may not be evident due to either the relatively young age of the studied individuals (such that meaningful neurodegeneration is not yet present) or due to compensation tissue strategies. In this regard, in our previous study we observed that structurally atrophic areas of the cortex in this group displayed increased metabolic activity, possibly as an early compensation strategy (28), which provides plausibility to this explanation.

Last, in line with several prior reports exploring the relationship between β-Amyloid deposition and depression in persons at risk of LOAD (64–67), we did not observe associations between regional PET-PiB signal and BDI scores in our sample of O-LOAD. However, the unexpected finding of a regional pattern of amyloid deposition in relation to self-reported symptoms of depression in middle-aged healthy controls, supports prior evidence on the role of distinct areas of the insular cortex (68,69), especially on the right hemisphere, as a major hub integrating interoceptive information in the posterior insula (70), with visceral regulation and encoding of emotional salience in the anterior insula (70,71). In this setting, impaired interoceptive integration by β-Amyloid in the posterior insula can result in dysregulation of this circuit and a higher report of depressive symptomatology. By contrast, increased β-Amyloid in the anterior insular cortices may impair emotional salience and thus result in a blunted subjective experience of negative affectivity. This is in line with prior findings in patient populations, describing reduced insular-anterior cingulate cortex connectivity with impaired salience processing and its behavioral correlate of emotional blunting (72). While compatible with existing knowledge on the regulation of negative emotion, we believe that this information should probably not be interpreted as a pathogenetic mechanism of clinical depression in middle age, for several reasons. Importantly, it needs to be emphasized that only a minor proportion of HC participants had BDI scores within a clinically meaningful range. In this setting, we speculate that rather than providing direct clues to mechanisms of MDD, regional β-Amyloid deposition influences limbic physiology in a nuanced way, and affects the subjective feeling of emotional status and visceral experience, thus contributing to how individual participants report these sensations in the self-administered BDI tool. In this scenario, rather than assigning a “protective” or antidepressant effect to deposition of abnormal pleated forms of β-Amyloid revealed by PET-PiB and deposited in bilateral anterior insulae, putamina, and left thalamus, our results point to the complex role of these structures (especially the right anterior insular cortex as the prime hub of emotional salience) in the experience of visceral sensations conducive to instant emotional status, which is then reported in this self-administered tool (73). Nonetheless, these results still support the role of anterior insular cortex and portions of the basal ganglia in the experience of negative emotion and unpleasantness, and as such, warrant further exploration of these structures as the use of anatomically precise deep brain neuromodulation techniques becomes more widespread (74,75).

There are several important limitations to take into account while interpreting the results of the present study. While deeply phenotyped by both MRI and PET imaging, the sample size of both O-LOAD and HC participants is relatively small, and there is a significant possibility that we may have overlooked significant imaging findings reflective of mechanisms of depressive symptoms in either group, including subtle PET-FDG changes resulting from very early neurodegeneration. The second major limitation of this study is that it is a cross-sectional observation, with no follow-up observations. A longitudinal follow-up design would allow to establish, depending upon progression to clinical LOAD, the predictive value of the different markers of depressive symptoms described herein. Until that observation is made in larger samples able to detect rates of conversion to clinical dementia, the pathophysiological meaning of the present findings remains speculative. Also, the sample of the present study belongs to a single site in a Latin American urban setting, and it was composed entirely of Caucasian Hispanic subjects. As such, the generalizability of symptoms remains limited as well. Last, while we attempted to exclude participants with significant white matter disease from vascular origin, we cannot rule out that this factor may have influenced some of the imaging results related to BDI scores, obscuring otherwise relevant relationships. The same can be said of depressive symptoms, which were assessed only with BDI and not a deep clinical interview or diagnostic schedule to rule out the presence of a major depressive disorder. Chronic exposure to specific serotonin reuptake inhibitors in six participants possibly introduced heterogeneity in the sample studied.

These limitations notwithstanding, the present study established distinct structural and functional neuroimaging correlates of depression in individuals with at least one parent with a well-established diagnosis of LOAD, and a comparable sample with no history of LOAD up to a 4th degree of kinship. While preliminary, these results offer a foundation for establishing heuristically valid hypotheses on the potential role of incipient neurodegenerative phenomena in the formation of depression symptoms after middle age, and as such, warrant further exploration on the mechanistic pathways of this disorder at different life stages.

## Acknowledgments

Effort on this article was supported by awards from NIMH (R01MH137331 and R21MH138911 to SMG) and the former Agencia de Promocion, Argentine government (PICT-2014-0633 to SMG and PICT-2019-02328 to MFV and SMG). The views expressed here are the authors’ and do not reflect the position or policies of the National Institutes of Health or the Argentine government.

